## Supplemental Figure 1 for "A Simple and Rapid Protocol for the Isolation of Murine Bone Marrow Suitable for the Differentiation of Dendritic Cells"

**Supplementary Figure 1**

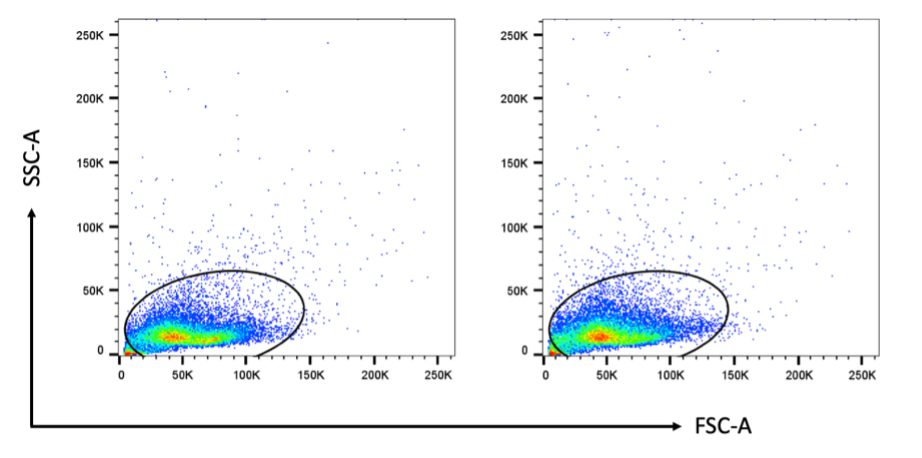

**A**

**B**

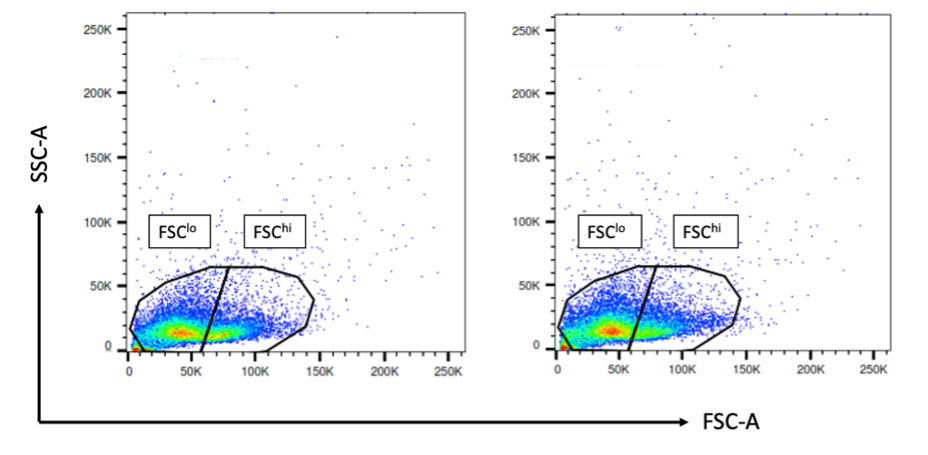

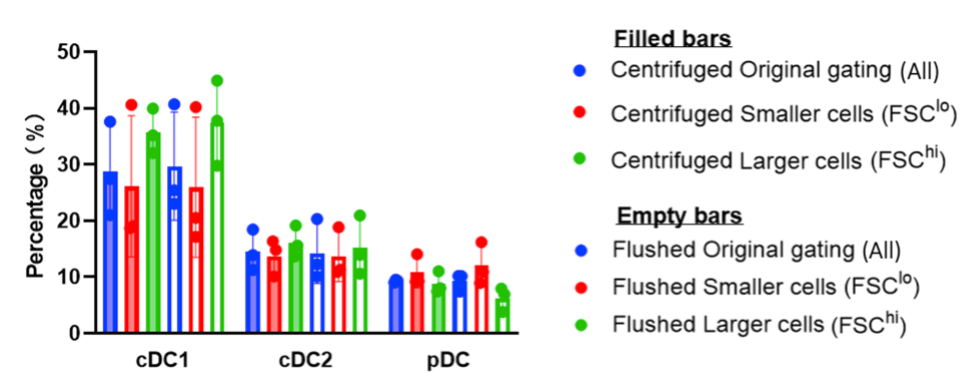

**C**

**The proportions of DC subsets in the culture based on cell size.** **(A)** The SSC-A/FSC-A gating including all cells in a wide gate for flushed cells (left) and centrifuged cells (right). A representative experimental day is shown. **(B)** Flushed cells (left) and centrifuged cells (right) from the same experimental day as (A), gated according to cell size (FSC-A). Cells smaller in size are gated as FSC^lo^ and larger cells as FSC^hi^. **(C)** The proportions of different DC subsets based on FSC-A gating, comparing all cells (original gating from A, blue), smaller cells (FSC^lo^, red), and larger cells (FSC^hi^, green), for both bone marrow isolation methods (centrifuged or flushed).
